# Evidence of causal effect of major depression on alcohol dependence: Findings from the Psychiatric Genomics Consortium

**DOI:** 10.1101/412098

**Authors:** Renato Polimanti, Roseann E. Peterson, Jue-Sheng Ong, Stuart MacGregor, Alexis C. Edwards, Toni-Kim Clarke, Josef Frank, Zachary Gerring, Nathan A. Gillespie, Penelope A. Lind, Hermine H. Maes, Nicholas G. Martin, Hamdi Mbarek, Sarah E. Medland, Fabian Streit, Major Depressive Disorder Working Group of the Psychiatric Genomics Consortium, Substance Use Disorder Working Group of the Psychiatric Genomics Consortium, 23andMe Research Team, Arpana Agrawal, Howard J. Edenberg, Kenneth S. Kendler, Cathryn M. Lewis, Patrick F. Sullivan, Naomi R. Wray, Joel Gelernter, Eske M. Derks

**Affiliations:** Department of Psychiatry, Yale University School of Medicine and VA CT Healthcare Center, West Haven, Connecticut, USA; Virginia Institute for Psychiatric and Behavioral Genetics, Department of Psychiatry, Virginia Commonwealth University, Richmond, Virginia, USA; Statistical Genetics, QIMR Berghofer, Brisbane, Australia; Division of Psychiatry, University of Edinburgh, Edinburgh, UK; Department of Genetic Epidemiology in Psychiatry, Central Institute of Mental Health, Medical Faculty Mannheim, Heidelberg University, Mannheim, Baden-Württemberg, Germany; QIMR Berghofer, Translational Neurogenomics, Brisbane, Australia; QIMR Berghofer, Psychiatric Genetics, Brisbane, Australia; Department of Human and Molecular Genetics, Massey Cancer Center, Virginia Commonwealth University, Richmond, Virginia, USA; QIMR Berghofer, Genetic Epidemiology, Brisbane, Australia; Department of Biological Psychology & EMGO+ Institute for Health and Care Research, Vrije Universiteit Amsterdam, Amsterdam, the Netherlands; 23andMe, Inc., Mountain View, California, USA; Department of Psychiatry, Washington University School of Medicine, Saint Louis, Missouri, USA; Department of Biochemistry and Molecular Biology, Indiana University School of Medicine, Indianapolis, Indiana, USA; Social, Genetic and Developmental Psychiatry Centre, Institute of Psychiatry, Psychology & Neuroscience, King’s College London, UK; Department of Medical Epidemiology and Biostatistics, Karolinska Institutet, Stockholm, Sweden; Department of Psychiatry, University of North Carolina at Chapel Hill, Chapel Hill, North Carolina, USA; Department of Genetics, University of North Carolina at Chapel Hill, Chapel Hill, North Carolina, USA; Institute for Molecular Bioscience, The University of Queensland, Brisbane, Australia; Queensland Brain Institute, The University of Queensland, Brisbane, Australia; Departments of Genetics and Neuroscience, Yale University School of Medicine, New Haven, Connecticut, USA

**Keywords:** major depression, alcohol dependence, alcohol consumption, genetic correlation, genome-wide association, Mendelian randomization

## Abstract

**Background:** Despite established clinical associations among major depression (MD), alcohol dependence (AD), and alcohol consumption (AC), the nature of the causal relationship between them is not completely understood.

**Methods:** This study was conducted using genome-wide data from the Psychiatric Genomics Consortium (MD: 135,458 cases and 344,901 controls; AD: 10,206 cases and 28,480 controls) and UK Biobank (AC-Frequency: from “daily or almost daily” to “never”, 438,308 individuals; AC-Quantity: total units of alcohol per week, 307,098 individuals). Linkage disequilibrium score regression and Mendelian Randomization (MR) analyses were applied to investigate shared genetic mechanisms (horizontal pleiotropy) and causal relationships (mediated pleiotropy) among these traits.

**Outcomes:** Positive genetic correlation was observed between MD and AD (rg_MD-AD_=+0.47, *P*=6.6×10^-10^). AC-Quantity showed positive genetic correlation with both AD (rg_AD-AC-Quantity_=+0.75, *P*=1.8×10^-14^) and MD (rg_MD-AC-Quantity_=+0.14, *P*=2.9×10^-7^), while there was negative correlation of AC-Frequency with MD (rg_MD-AC-Frequency_=-0.17, *P*=1.5×10^-10^) and a non-significant result with AD. MR analyses confirmed the presence of pleiotropy among these traits. However, the MD-AD results reflect a mediated-pleiotropy mechanism (i.e., causal relationship) with a causal role of MD on AD (beta=0.28, *P*=1.29×10^-6^) that does not appear to be biased by confounding such as horizontal pleiotropy. No evidence of reverse causation was observed as the AD genetic instrument did not show a causal effect on MD.

**Interpretation:** Results support a causal role for MD on AD based on genetic datasets including thousands of individuals. Understanding mechanisms underlying MD-AD comorbidity not only addresses important public health concerns but also has the potential to facilitate prevention and intervention efforts.

**Funding:** National Institute of Mental Health and National Institute on Drug Abuse.

**Putting data into context:** *Evidence before this study:* We searched PubMed up to August 24, 2018, for research studies that investigated causality among alcohol-and depression related phenotypes using Mendelian randomization approaches. We used the search terms “alcohol” AND “depression” AND “Mendelian Randomization”. No restrictions were applied to language, date, or article type. Ten articles were retrieved, but only two were focused on alcohol consumption and depression-related traits. The studies were based on genetic variants in *alcohol dehydrogenase* (*ADH*) genes only, did not find evidence for a causal effect of alcohol consumption on depression phenotypes, with one study finding a causal effect of alcohol consumption on alcoholism. Both studies noted that future studies are needed with increased sample sizes and clinically derived phenotypes. To our knowledge, no previous study has applied two-sample Mendelian randomization to investigate causal relationships between alcohol dependence and major depression. Twin studies show genetic factors influence susceptibility to MD, AD, and alcohol consumption. Differently from observational approaches where several studies have investigated the relationship between alcohol-and depression-related phenotypes, very limited use of molecular genetic data has been applied to investigate this issue. Additionally, the use of genetic information has been shown to be less biased by confounders and reverse causation than observation data. However, genetic approaches, like Mendelian randomization, require large sample sizes to be informative.

*Added value of this study:* In this study, we used genome-wide data from the Psychiatric Genomic Consortium and UK Biobank, which include information regarding hundred thousands of individuals, to test the presence of shared genetic mechanisms and causal relationships among major depression, alcohol dependence, and alcohol consumption. The results support a causal influence of MD on AD, while alcohol consumption showed shared genetic mechanisms with respect to both major depression and alcohol dependence.

*Implications of all the available evidence:* Given the significant morbidity and mortality associated with MD, AD, and the comorbid condition, understanding mechanisms underlying these associations not only address important public health concerns but also has the potential to facilitate prevention and intervention efforts.

## INTRODUCTION

Major depression (MD) and alcohol dependence (AD) are common psychiatric disorders, contribute substantially to global morbidity, and often co-occur.^1,2^ Epidemiological studies report that individuals with MD are at increased risk for AD and vice versa.^3^–^5^ Leading hypotheses suggest these associations may be due to shared risk factors (genetic and environmental) or causal processes of one disorder leading to the other, such as the self-medication hypothesis of MD.^6^ However, the mechanisms underlying MD-AD dual diagnosis remain unclear.

Twin studies show genetic factors influence susceptibility to MD, AD, and alcohol consumption.^7^–^9^ Large-scale genome-wide association studies (GWAS) have identified risk variants for these disorders and have revealed polygenic architectures with multiple common variants.^10^–^14^ Twin studies report moderate shared genetic liability between MD and AD, estimating the genetic correlation from 0.3 to 0.6.^15,16^ Although emerging molecular genetic studies have reported shared genetic risk between these disorders, they have not yet illuminated mechanisms of association underlying genetic correlations.^11^–^13,17^–^19^ That is, questions remain whether these traits show genetic correlation because of shared genetic effects independently on each trait (i.e., horizontal pleiotropy)^20^ or because of causal processes (e.g., mediated pleiotropy).

GWAS data can be used to assess causal mechanisms by applying Mendelian randomization (MR). MR is an instrumental variables technique that uses genetic variants to index if an observational association between a risk factor (e.g., MD) and an outcome (e.g., AD) is consistent with a causal effect (e.g., MD causes AD). MR relies on random assortment of genetic variants during meiosis which are typically unassociated with confounders since they are randomly distributed in the population at birth. The differences in outcome between those who carry genetic risk variants and those who do not can be attributed to the difference in the risk factor. The validity of the genetic instrument is dependent on meeting three core assumptions: (1) the genetic variant is associated with the risk factor/exposure; (2) the genetic variant is not associated with confounders; and (3) the genetic variant influences the outcome only through the risk factor. Although random controlled trials (RCTs) are considered the gold standard for establishing causality, MR is a viable alternative to provide support for causal mechanisms, especially when RCTs are not possible or ethical.

Here, we leverage GWAS summary statistics generated by large datasets from the Psychiatric Genomics Consortium (PGC) and the UK Biobank to estimate genetic correlations between MD, AD, and two measures of alcohol consumption (AC: AC-Quantity, AC-Frequency) via linkage disequilibrium (LD) score regression.^17,21^ Further, we investigated support for causal mechanisms linking these psychiatric disorders and alcohol consumption via two-sample Mendelian randomization analyses, which use genetic variants to assess whether an exposure has a causal effect on an outcome in a non-experimental setting.^22^

## METHODS

### Samples

#### 1. Major depression (MD)^11^

MD summary association data were obtained from the latest GWAS meta-analysis including 135,458 MD cases and 344,901 controls from the MD working group of the PGC (PGC-MDD2), which included 7 cohorts: (1) 29 samples of European-ancestry with MD-cases required to meet international consensus criteria (DSM-IV, ICD-9, or ICD-10) for a lifetime diagnosis of MD established using structured diagnostic instruments from assessments by trained interviewers, clinician-administered checklists, or medical record review and controls in most samples were screened for the absence of lifetime MD (22/29 samples), and randomly selected from the population; (2) Generation Scotland employed direct interviews; (3) iPSYCH (Denmark) used national treatment registers; (4) deCODE (Iceland) used national treatment registers and direct interviews; (5) GERA used Kaiser-Permanente treatment records (CA, US); (6) UK Biobank combined self-reported MD symptoms and/or treatment for MD by a medical professional; and (7) 23andMe used self-report of treatment for MD by a medical professional. All controls included in datasets 2-7 were screened for the absence of MD.

#### 2. Alcohol Dependence (AD)^12^

AD summary association data from unrelated subjects of European descent (10,206 cases; 28,480 controls) were obtained from GWAS meta-analysis of 14 cohorts conducted by the PGC Substance Use Disorder Workgroup. Detailed descriptions of the AD samples have been previously reported.^12^ In brief, AD was defined as meeting criteria for a DSM-IV (or DSM-IIIR in one instance) diagnosis of AD and with the exception of three cohorts with population-based controls (*n*=7,015), all controls were screened for AD. Individuals with no history of drinking alcohol and those meeting criteria for DSM-IV alcohol abuse were additionally excluded as controls where applicable (i.e., where these data were available).

#### 3. UK Biobank (UKB) - alcohol consumption quantity and frequency

The UK Biobank cohort consists of 502,000 middle-aged (40-69 years) individuals recruited from the United Kingdom. The UK Biobank records extensive (*n*>2000) phenotypes of the participants ranging from anthropometric traits, to disease status, to lifestyle behaviours.

Information on alcohol intake was obtained through various self-report questionnaires. Frequency of consumption (AC-Frequency) was assessed in 501,718 participants (UKB field IDs: 1558) with the item “About how often do you drink alcohol?”. Frequency was originally assessed at a scale ranging from 1 (daily or almost daily) to 6 (never), but was reverse coded so that a lower score represented less frequent drinking. Figure S3 shows the distribution in the UKB population. In those who drink at least once or twice a week, information on quantity of consumption (AC-Quantity) was assessed (*n*=348,039). AC-Quantity was assessed based on the average weekly alcohol intake for five general classes: red wine (field ID: 1568), champagne plus white wine (field ID: 1578), spirits (field ID: 1598), beer plus cider intake (field ID: 1558), and fortified wine (field ID: 1608). The following item was used: “In an average WEEK, how many servings of {class of alcohol} would you drink?”. More detailed information on quality checks can be found in the Supplemental Methods.

For a complete description of the UKB genotype curation, please see the report by Ong and colleagues.^23^ All participants provided informed written consent, the study was approved by the National Research Ethics Service Committee North West – Haydock, and all study procedures were performed in accordance with the World Medical Association Declaration of Helsinki ethical principles for medical research. In brief, approximately 488,000 participants were genotyped on custom-designed Affymetrix UK BiLEVE Axiom or UK Biobank Axiom arrays (Affymetrix Santa Clara, USA), which produced a combined total of 805,426 markers in the released data. Following standard quality control (QC) the dataset was phased and ∼96M genotypes were imputed using Haplotype Reference Consortium (HRC) and UK10K haplotype resources.^24^–^26^ Due to the UKB’s reported QC issues with non-HRC SNPs, we retained only the ∼40M HRC SNPs for analysis. In light of a large number of related individuals in the UKB cohort, the GWAS was performed using BOLT-LMM which is a linear mixed model framework that explicitly models the genetic relatedness within the sample.^27^

Among the 487,409 individuals who passed initial genotyping QC, 409,694 participants had white-British ancestry, based on self-reported ethnicity and genetic principal components. To maximize our effective sample size, we also included UKB participants if their self-reported ancestry was not white-British (this includes a substantial number of individuals reporting their ancestry as “Irish” or “any other white background”) but their first two genetic principal components fell within the region of those that are classified white-British in the *n* = 409,694 set. Using these criteria, we identified 438,870 individuals for this study who are genetically similar to those of white-British ancestry. After exclusion of ethnic outliers, we included 438,308 participants in the AC-Frequency and 307,098 participants in the AC-Quantity GWAS.

### Sample Overlap

Among the samples included in the MD and AD GWAS, three cohorts (of 22 AD cohorts and 35 MD cohorts) were present in both analyses and some individuals from these cohorts may overlap. LD score regression is not biased by sample overlap.^21^ Simulations on two-sample MR methods demonstrated that the relative bias (which may be toward a null direction) with 50% sample overlap was 5% and with 30% sample overlap was 3%.^28^ To quantify potential bias from GWAS summary statistics due to overlapping samples, we used LambdaMeta implemented in GEnetic Analysis Repository (GEAR).^29^ Under the null hypothesis, LambdaMeta is 1 when no “sample-overlap” effect is affecting the pair of summary statistics. When the summary statistics are affected by sample overlap, LambdaMeta < 1; when there are technical differences, LambdaMeta > 1. LambdaMeta between MD and AD GWAS was 1.0021, suggesting results should not be significantly biased due to sample overlap.

### SNP-based heritability analysis

The proportion of variance in phenotypic liability that could be explained by the aggregated effect of all SNPs (h^2-^SNP) was estimated using LD-Score Regression analysis (see Supplemental Methods).^21^ For this analysis, we included in the regression 1,217,311 SNPs that were present in the HapMap 3 reference panel. Analyses were performed using pre-computed LD scores based on 1000 Genomes Project reference data on individuals of European ancestry (available for download at https://data.broadinstitute.org/alkesgroup/LDSCORE/). The h^2^-SNP estimates for the two binary traits were converted to the liability scale, using sample prevalence of 0.159 for AD and 0.15 for MD. It should be noted that h^2^-SNP estimates may be slightly underestimated since summary statistics were derived from a linear mixed model analysis (BOLT-LMM) and mixed models may change the expected behavior of the mean chi-square.

### Genetic correlations between MD, AD, AC-Quantity, and AC-Frequency

We used cross-trait LD-Score regression to estimate the bivariate genetic correlations between MD, AD, and AC using GWA summary statistics.^17^ For each pair of traits, the genetic covariance is estimated using the slope from the regression of the product of z-scores from two GWA studies on the LD score. The estimate represents the genetic covariation between the two traits based on all polygenic effects captured by SNPs. To correct for multiple testing, we adopted a Bonferroni corrected *P*-value threshold of significance of 0.05/6=0.0083. Analyses were performed using pre-computed LD scores based on 1000 Genomes Project reference data on individuals of European ancestry.

### Mendelian Randomization

To assess causality among MD, AD, AC-Quantity, and AC-Frequency, we used GWAS summary association data to conduct two-sample MR analyses.^17,22^ Since different MR methods have sensitivities to different potential issues, accommodate different scenarios, and vary in their statistical efficiency,^30^ we considered multiple MR methods (Table S2). These include methods based on median,^30,31^ mean,^32^ and mode;^33^ and various adjustments, such as fixed vs. random effects,^31^ Rucker framework,^31,34^ and Steiger filtering.^35^ We verified the stability of the results, comparing the effect directions across the different MR-variant filtering methods (Table S2).MR-Egger regression intercept was considered to verify the presence of pleiotropic effects of the SNPs on the outcome (i.e., to verify whether the instrumental variable is associated with the outcome independently from its association with the exposure).^36^ In total we performed 17 MR tests (Table S1). This number is due to the fact we were not able to test AD using a genetic instrument based on genome-wide significant loci and, since we are conducting a two-sample MR analysis, we did not test causal relationship between AC-Quantity and AC-Frequency because they are based on UK Biobank cohort. For the variants included in the instrumental variable, we performed linkage disequilibrium clumping by excluding alleles that have R^2^≥0.01 with another variant with a smaller association *P*-value considering a 1Mb window.

Additionally, during the harmonization of exposure and outcome data, palindromic variants with an ambiguous allele frequency (i.e., minor allele frequency close to 50%) were removed from the analysis to avoid possible issues.^36,37^ The variants included in each genetic instrument used in the present analysis are listed in Table S6. For each exposure, two instrumental variables were built considering genome-wide significant (GWS) loci (P<5×10^-8^) and suggestive loci (P<5×10^-5^). To ensure the reliability of the significant findings, we performed heterogeneity tests based on three different methods: inverse-variance weighted, MR-Egger regression, and maximum likelihood (Table S5). To further confirm the absence of possible distortions due to heterogeneity and pleiotropy, we tested the presence of horizontal pleiotropy among the variants included in the genetic instrument using MR-PRESSO.^38^ Finally, funnel plot and leave-one-out analysis were conducted to identify potential outliers among the variants included in the genetic instruments tested. The MR analyses were conducted using the TwoSampleMR R package.^39^

## RESULTS

### SNP-based heritabilities and genetic correlations

We confirmed previously reported heritability estimates of MD (h^2^-SNP=8.5%, SE=0.003, *K*=0.15) and AD (h^2^-SNP = 9.0%, SE=0.019, *K*=0.16), with K defined as the disease prevalence in the population.^11,12^ The h^2^-SNP of AC-Frequency, which has not been previously reported, was estimated at 8.0% (SE=0.003). The h^2^-SNP of AC-Quantity using LD-score regression was estimated at 6.9% (SE=0.004), which is lower than the GCTA-estimate reported by Clarke et al. (13%) who analyzed a smaller subset (n=112,117)^13^ from the current data (n=307,098). The lower estimate may be explained by differential methodology (i.e., LD-score regression vs.GCTA) and by the fact that the first release of UKB included a subset of individuals that was selected based on smoking and may be less representative of the general population than the current sample.^40^ These h^2^-SNP estimates are capturing 17-23% of heritabilities reported by twin studies.^7,9^

The high genetic correlation between AD and AC-Quantity (rg_AD-AC-Quantity_=+0.75, 95%CI=[0.56, 0.94], *P*=1.8×10^-14^) (Figure 1) suggests that these phenotypes capture overlapping constructs and that quantity of consumption is an indicator of problematic alcohol use. Of note, the genetic correlation between AD and AC-Frequency is not significantly different from zero, indicating that it is not a reliable indicator of genetic risk for AD.

**Figure 1:**
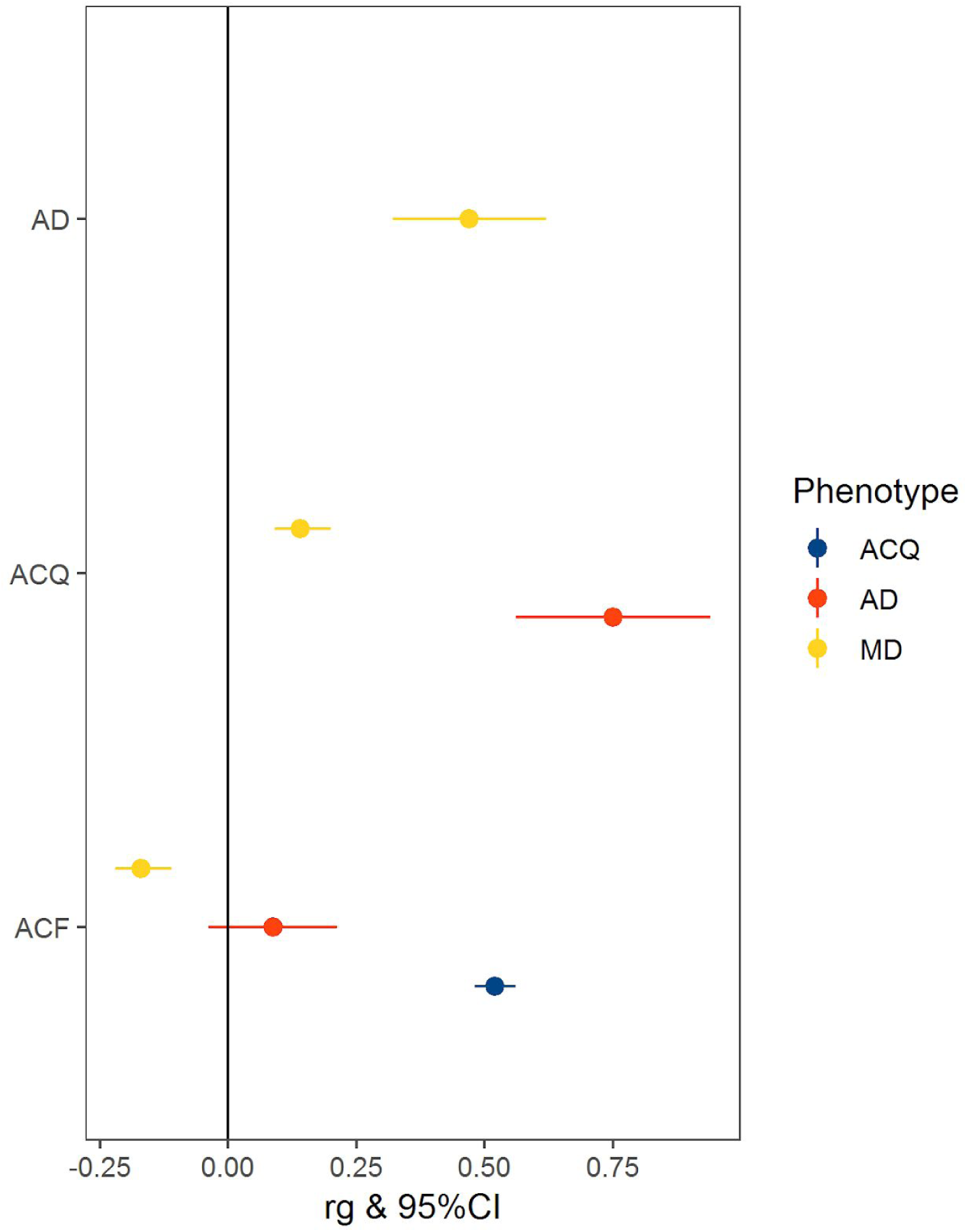
Genetic correlations of Major depression (MD), Alcohol Dependence (AD), and alcohol consumption quantity (ACQ) and frequency (ACF) Note: AD-Alcohol Dependence; MD-Major depression; ACQ-quantity of alcohol consumption; ACF-frequency of alcohol consumption.

Consistent with twin studies, MD and AD show moderate overlap of genetic factors (rg_MD-AD_=+0.47, 95%CI=[0.32,0.62], *P*=6.6×10^-10^).^15,16^ A significant genetic correlation between AC-Quantity and AC-Frequency was observed (rg_AC-Quantity-AC-Frequency_=+0.52, 95%CI=[0.48, 0.56], *P*=1.3×10^-149^), but MD showed significant correlations with these traits in opposite directions (rg_MD-AC-Quantity_=+0.14, 95%CI=[0.09, 0.20], *P*=2.9×10^-7^; rg_MD-AC-Frequency_=-0.17, 95%CI=[-0.22, −0.11], *P*=1.5×10^-10^).

### Mendelian Randomization

SNP-based heritability and genetic-correlation results demonstrate that our GWAS data are informative with respect to the phenotypic variance explained by common genetic variation and shared genetic factors among the phenotypes investigated. However, genetic correlation analyses do not distinguish between mediated pleiotropy (i.e., genetic variants exert an effect on one trait through their association with another one) and horizontal pleiotropy (i.e., genetic variants independently affect multiple traits). Therefore, we investigated the presence of mediated pleiotropy via two-sample MR as this allows us to test for causative mechanisms linking MD, AD, AC-Frequency, and AC-Quantity. This is a strategy to investigate causal relationships in which evidence on the associations of genetic variants (i.e., instrumental variable) with the risk factor (i.e., exposure) and with the outcome are derived from two samples.^22,41^ The instrumental variables were built considering GWS loci (*P*<5×10^-8^) and suggestive loci (*P*<5×10^-5^). Since different MR methods have sensitivities to different potential issues, we considered 28 MR/variant-filtering approaches (Table S2). A Bonferroni correction (*P*<1.05×10^-4^) was applied to correct for the number of MR tests performed (*n*=17; Table S1) and the number of methods/variant-filtering considered for each test (*n*=28; Table S2). Of the 17 MR tests conducted, we observed that 14 survived multiple testing correction (Table 1). This outcome was expected due to the strong genetic correlations observed among the traits investigated. To verify that the significant results were not due to the presence of biases in the genetic instruments, we conducted three main sensitivity analyses: i) inspected consistency of direction of effects across MR methods (Table S3); ii) tests of horizontal pleiotropy between the exposure and the outcome (MR-Egger regression intercept *P*>0.1; Table S4); iii) assessed heterogeneity of effect sizes among the variants included in the genetic instrument (heterogeneity test *P*>0.05; Table S5). Of 14 MR tests surviving Bonferroni multiple testing correction, only the causal relationship of MD on AD passed all three sensitivity analyses. We observed that the MD instrumental variable based on suggestive variants (259 SNPs) was associated with AD (fixed-effect inverse-variance weighted method: Beta=0.28, *P*=1.3×10^-6^; Figure 2). A similar effect size was also observed for the MD instrumental variable based on GWS loci (40 SNPs; fixed-effect inverse-variance weighted method: Beta=0.27, *P*=0.054). Results indicated that MD is associated with a 32% increase in the odds for AD risk per unit increase in the log(OR) for MD (95%CI: 18-48%) and were consistent across multiple MR approaches (Table S3.17). As mentioned above, the MD genetic instrument did not show evidence of horizontal pleiotropic effects as demonstrated by MR-Egger regression intercept (*P*=0.297, Table S4), confirming that the causal effect of MD on AD does not appear to be biased by horizontal pleiotropy. The heterogeneity tests indicated no evidence of heterogeneity in the MD-AD result (*P*>0.13; Table S5). The MR-PRESSO global test^38^ also supported the absence of horizontal pleiotropy (RSSobs=285.6, *P*=0.143). Finally, the funnel plot and leave-one-out analyses provided supplementary support that the MD-AD result was not biased by outliers included in the genetic instrument (Figure S1). The same MD genetic instrument also showed significant effects on AC-Quantity and AC-Frequency (Table 1), but, in contrast to the AD outcome, these causal effects showed evidence of non-consistency across MR methods, heterogeneity, and horizontal pleiotropy (Table S3-S5). No reverse causal effect was observed between AD genetic instrument and MD (fixed-effect inverse-variance weighted method: Beta=0.01, *P*=0.1), which also showed non-concordant direction of effects across MR methods (Figure S2). Conversely, the AD genetic instrument showed significant effects on AC-Quantity and AC-Frequency but were affected by heterogeneity and horizontal pleiotropy (Table 1; Table S4-S5).

**Table 1:** Results of the most significant MR approach among those surviving Bonferroni multiple testing correction for each of the MR tests conducted. All top-results reported in the table were obtained using fixed effects and tophits adjustments (see Table S2).

| Test | Method | SNP<br>*n* | Estimate | *P* | Concordance | Het.<br>(*P*>0.05) | MR-Egger<br>Intercept<br>(*P*>0.1) |
| --- | --- | --- | --- | --- | --- | --- | --- |
| ACF→AD | Egger | 92 | -1.97 | 2.51E-11 | pass | violated | violated |
| ACFe-5→AD | Egger | 773 | -0.75 | 3.24E-06 | pass | violated | violated |
| ACFe-5→MD | IVW | 795 | 0.07 | 6.50E-20 | violated | violated | violated |
| ACQ→AD | IVW | 31 | 0.12 | 3.81E-11 | pass | violated | violated |
| ACQ→MD | IVW | 30 | 0.01 | 8.06E-05 | violated | violated | pass |
| ACQe-5→AD | IVW | 385 | 0.06 | 7.34E-20 | pass | violated | violated |
| ACQe-5→MD | IVW | 405 | 0.01 | 2.43E-17 | violated | violated | violated |
| ADe-5→ACF | Egger | 96 | -0.05 | 9.11E-48 | pass | violated | violated |
| ADe-5→ACQ | IVW | 95 | 0.26 | 3.26E-36 | pass | violated | pass |
| MD→ACF | IVW | 36 | 0.05 | 4.06E-08 | pass | violated | pass |
| MD→ACQ | Egger | 36 | -4.77 | 1.93E-10 | pass | violated | violated |
| MDe-5→ACF | IVW | 252 | 0.02 | 1.27E-08 | violated | violated | pass |
| MDe-5→ACQ | IVW | 251 | 0.31 | 1.46E-06 | violated | violated | violated |
| MDe-5→AD | IVW | 259 | 0.28 | 1.29E-06 | pass | pass | pass |

Note: ACF, alcohol consumption frequency; ACQ, alcohol consumption quantity; AD, alcohol dependence; MD, major depression; Het., heterogeneity test; suggestive loci ( $P < 5 \times 10^{-5}$ ).

**Figure 2:**
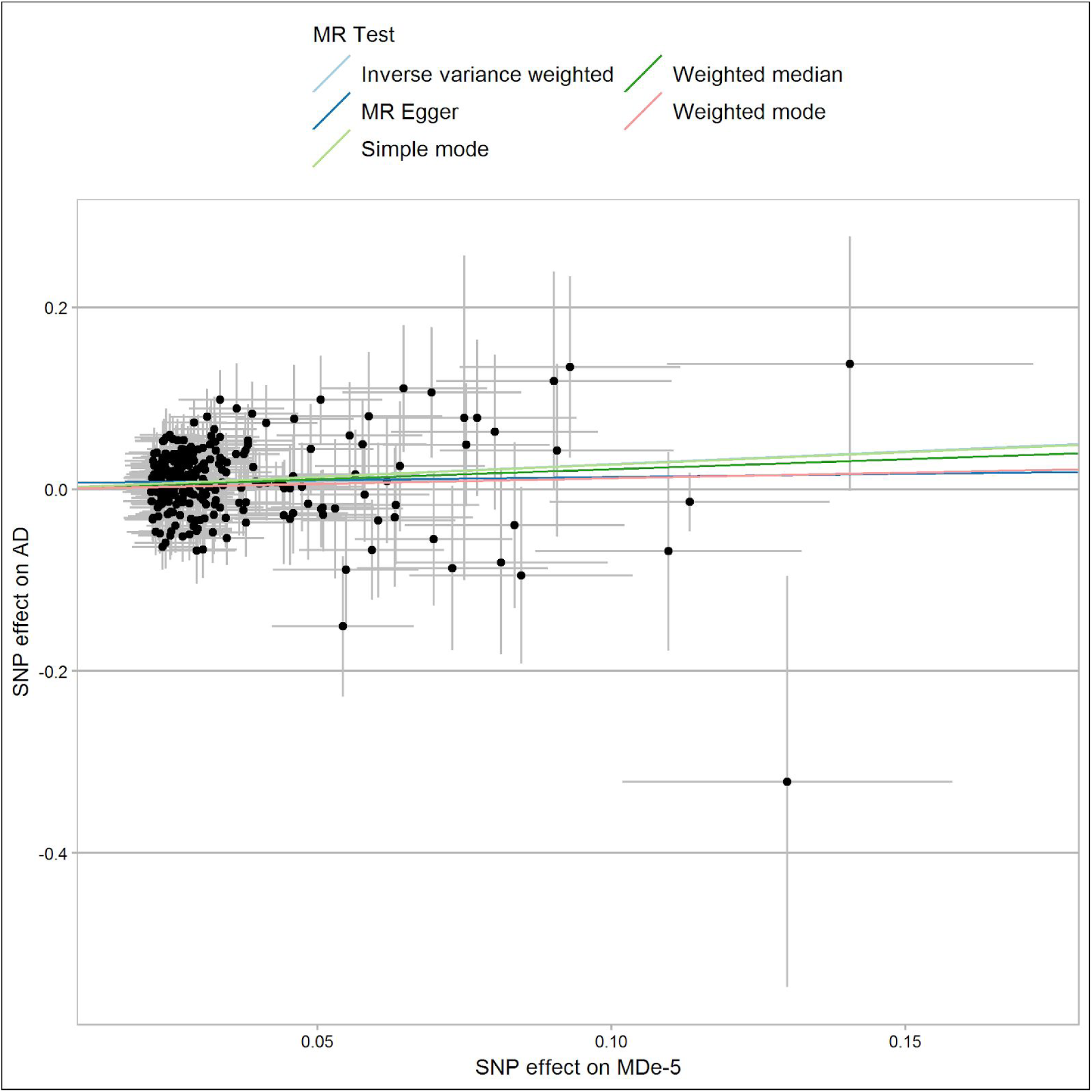
SNP-exposure (MD associations, logOR) and SNP-outcome (AD associations, logOR) coefficients used in the MR analysis. Error bars (95% CIs) are reported for each association.

## DISCUSSION

Data from large-scale GWAS are redefining the boundaries of psychiatric disorders, identifying the contribution of common risk alleles and pervasive genetic correlations. Here, leveraging the polygenic architecture of these complex traits and the large sample size of GWAS results from the Psychiatric Genomics Consortium and UK Biobank, we observed genetic overlap between MD and AD and provide support for a causal effect of MD on AD, which does not appear to be affected by horizontal pleiotropy or other detectable biases. To our knowledge, this is the first report of causality between MD and AD based on molecular genetic information. Consistent with the two previously published MR studies,^18,19^ we did not find a robust causal association of alcohol consumption on depression.

We detected significant genetic correlations between all pairs of phenotypes, except for AD and AC-Frequency, suggesting that frequency of alcohol consumption is not a good proxy for alcohol dependence. In contrast, the genetic correlation between AD and AC-Quantity is high at rg=0.75. Of note, our estimate is higher than previously reported by Walters et al. (rg=0.37) who used the same GWAS results for AD^12^ but the earlier, smaller subset of the UKB data on consumption.^13^ However, our estimate is comparable to the estimate reported by Walters et al.(rg_AD-AC-Quantity_=+0.70) who calculated the genetic correlation between Alcohol Dependence and alcohol consumption from the Alcohol Genome-wide Association and Cohorts for Aging and Research in Genomic Epidemiology Plus consortium,^14^ suggesting that the genetic correlation between these traits is indeed high.^12^ Therefore, AC-Quantity, which can be collected at low cost in large samples, can potentially be used as a proxy for Alcohol Dependence. This is especially important since the sample size of the Alcohol Dependence GWAS is relatively small resulting in few genome-wide significant associations.

The finding of a negative correlation between AC-Frequency and MD may seem counterintuitive, but is supported by earlier studies that report opposing effects of quantity and frequency on health. For example, high drinking quantity is associated with increased all-cause mortality risk while frequency of drinking does not show such an association.^42^ Additionally, in line with our findings, it was recently shown that MD is associated in opposite directions with two aspects of problematic drinking as assessed by the AUDIT: AUDIT-consumption (i.e., assessing frequency of consumption) (rg=-0.23) and AUDIT-problematic consequences (rg=+0.26).^43^ Our findings support the notion that MD is genetically positively correlated with measures of problematic drinking (i.e., AD and AC-Quantity), but is negatively correlated with frequency of consumption.

In contrast to some epidemiological reports,^5^ our results do not support evidence of reverse causation, that is, AD causing MD. One could posit that this is due to the relative power of the AD instrumental variable compared to those for MD and AC given the greater number of genome-wide significant variants detected for those traits. However, we would like to argue that the null ADe-5→MD result could be due to the absence of a causal effect of AD on MD rather than a lack of power of the AD dataset for three reasons. Indeed, the AD genetic instrument showed different associations between the traits tested: significant causal effect with respect to alcohol consumption scales (ADe-5→ACF and ADe-5→ACQ), while no effect on MD (ADe-5→MD). The genetic-correlation results indicated that AD is informative of the pleiotropy (mediated or horizontal) with ACQ and MD. Additional support to the difference of AD with respect to ACQ and ACF is provided by the results of the sensitivity analyses conducted with respect to the MR tests based on the MDe-5 genetic instrument (MDe-5→AD, MDe-5→ACQ, and MDe-5→ACF). The absence of heterogeneity/pleiotropy in the “MDe-5→AD” causal effect and the presence of heterogeneity/pleiotropy in other MR tests conducted using the same genetic instrument (i.e., MDe-5→ACQ and MDe-5→ACF) are due to the different architectures of the alcohol-related traits investigated rather than the reduced power of AD dataset. Although the AD GWAS has a smaller sample size than the other GWAS used in the present analysis, it is informative of AD polygenic architecture as indicated by quantifiable and statistically significant SNP-based heritability and genetic correlation results. In particular, MD showed a much stronger genetic correlation with AD than that observed with alcohol consumption scales. However, we note that larger AD and MD datasets will be required to confirm the current findings using genetic instruments based on genetic variants that reached the more conservative genome-wide significance threshold.

In conclusion, these results support the utility of using genetic approaches to advance the understanding of complex trait comorbidities. Given the significant morbidity and mortality associated with MD, AD, and the comorbid condition, understanding mechanisms underlying these associations not only address important public health concerns but also has the potential to facilitate prevention and intervention efforts. As discovery GWAS increase in sample size, future research will have the power to examine patterns of genetic correlation and causal mechanisms by important stratifications such as across diverse ancestries and sex.

## Acknowledgements

The Psychiatric Genomics Consortium (PGC): We are deeply indebted to the investigators who comprise the PGC, and to the hundreds of thousands of subjects who have shared their life experiences with PGC investigators. The PGC has received major funding from the National Institute of Mental Health and the National Institute on Drug Abuse (PGC3: U01 MH109528 and U01 MH109532, PGC2: U01 MH094421, PGC1:U01MH085520). The Substance Use Disorders Working Group of the Psychiatric Genomics Consortium (PGC-SUD) is supported by funds from NIDA and NIMH to MH109532 and, previously, with analyst support from NIAAA to U01AA008401 (COGA). We gratefully acknowledge the contributing studies and the participants in those studies without whom this effort would not be possible. For a full list of acknowledgements of all individual cohorts included in the PGC-SUD and PGC-MD groups, please see the original publications. Statistical analyses were carried out on the Genetic Cluster Computer (http://www.geneticcluster.org) hosted by SURFsara, which is financially supported by the Netherlands Scientific Organization (NWO 480-05-003) along with a supplement from the Dutch Brain Foundation and the VU University Amsterdam. Renato Polimanti was supported by a Young Investigator Grant from the American Foundation for Suicide Prevention. Roseann E. Peterson was supported by National Institutes of Health K01 grant MH113848. Nathan A. Gillespie was supported by National Institutes of Health R00 grant R00DA023549. This paper represents independent research part-funded by the National Institute for Health Research (NIHR) Biomedical Research Centre at South London and Maudsley NHS Foundation Trust and King’s College London. The views expressed are those of the authors and not necessarily those of the NHS, the NIHR or the Department of Health and Social Care. This work was conducted using the UK Biobank Resource (application number 25331). Collaborators for the 23andMe Research Team are: Michelle Agee, Babak Alipanahi, Adam Auton, Robert K. Bell, Katarzyna Bryc, Sarah L. Elson, Pierre Fontanillas, Nicholas A. Furlotte, David A. Hinds, Karen E. Huber, Aaron Kleinman, Nadia K. Litterman, Matthew H. McIntyre, Joanna L. Mountain, Elizabeth S. Noblin, Carrie A.M. Northover, Steven J. Pitts, J. Fah Sathirapongsasuti, Olga V. Sazonova, Janie F. Shelton, Suyash Shringarpure, Chao Tian, Joyce Y. Tung, Vladimir Vacic, Catherine H. Wilson. We thank the research participants of 23andMe.

## Authors’ Contributions

R.P., R.E.P., E.M.D. Designed this study and co-wrote the manuscript; R.P., J-S. O., E.M.D. Performed Statistical analyses; All other authors: Drafted or provided critical revision of the article and provided final approval of the version to publish.

## Conflicts of Interest

Members of the 23andMe Research Team are employees of 23andMe, Inc. and hold stock or stock options in 23andMe. All remaining authors declare no conflict of interest.

## Data availability statement

### Major Depression GWAS

The PGC’s policy is to make genome-wide summary results public. Summary statistics for a combined meta-analysis of PGC29 with five of the six expanded samples (deCODE, Generation Scotland, GERA, iPSYCH, and UK Biobank) are available on the PGC web site (https://www.med.unc.edu/pgc/results-and-downloads). Results for 10,000 SNPs for all seven cohorts are also available on the PGC web site. GWA summary statistics for the Hyde et al. cohort (23andMe, Inc.) must be obtained separately. These can be obtained by qualified researchers under an agreement with 23andMe that protects the privacy of the 23andMe participants. Contact David Hinds to apply for access to the data.

Researchers who have the 23andMe summary statistics can readily recreate our results by meta-analyzing the six cohort results file with the Hyde et al. results file from 23andMe.

### Alcohol Dependence GWAS

The PGC’s policy is to make genome-wide summary results public. Summary statistics will be made publicly available before publication of this article.

### Alcohol Consumption Quantity and Frequency GWAS

Summary statistics will be made publicly available through LD hub http://ldsc.broadinstitute.org/ldhub/beforepublication of this manuscript or can be obtained upon request from the corresponding author.

